# Biophysical Kv channel alterations dampen excitability of cortical PV interneurons and contribute to network hyperexcitability in early Alzheimer’s

**DOI:** 10.1101/2021.10.25.465789

**Authors:** Viktor Janos Oláh, Annie M Goettemoeller, Jordane Dimidschstein, Matthew JM Rowan

## Abstract

In Alzheimer’s disease (AD), a multitude of genetic risk factors and early biomarkers are known. Nevertheless, the causal factors responsible for initiating cognitive decline in AD remain controversial. Toxic plaques and tangles correlate with progressive neuropathology, yet disruptions in circuit activity emerge before their deposition in AD models and patients. Parvalbumin (PV) interneurons are potential candidates for dysregulating cortical excitability, as they display altered AP firing before neighboring excitatory neurons in prodromal AD. Here we report a novel mechanism responsible for PV hypoexcitability in young adult familial AD mice. We found that biophysical modulation of K^+^ channels, but not changes in mRNA expression, are responsible for dampened excitability. These K^+^ conductances could efficiently regulate near-threshold AP firing, resulting in gamma-frequency specific network hyperexcitability. Our findings suggest that posttranslational modulation of ion channels can reshape cortical network activity prior to changes in their gene expression in early AD.

## Introduction

Unraveling mechanisms that initiate cognitive decline in Alzheimer’s disease (AD) is a central aim in neuroscience. A prevailing model of AD posits that progressive deposition of toxic proteins spark a neuropathological cascade. However, recent work suggests that early cognitive dysfunction is uncoupled from these aggregates (Arroyo-García et al., 2021; Nuriel et al., 2017; Shimojo et al., 2020). Several alternative models for early cognitive decline are under consideration (De Strooper and Karran, 2016) including abnormal circuit activity (Busche and Konnerth, 2015; Busche et al., 2008; Cirrito et al., 2005; Davis et al., 2014; Pooler et al., 2013; Wu et al., 2016). Circuit hyperexcitability is evident in several mouse models of familial (FAD) and sporadic AD (Lamoureux et al., 2021; Minkeviciene et al., 2009; Nuriel et al., 2017) including at prodromal stages (Bai et al., 2017; Busche and Konnerth, 2015). Furthermore, abnormal brain activity is apparent in humans with mild cognitive impairment (Dickerson et al., 2005; Hämäläinen et al., 2007; Miller et al., 2008; Sperling et al., 2010) and in early FAD (Quiroz et al., 2010; Sepulveda-Falla et al., 2012). These shifts in circuit activity may result from dysfunctional neuronal firing and neurotransmission (Chen et al., 2018; Palop and Mucke, 2016). However, the cellular and molecular mechanisms underlying these neuronal deficits are not yet fully understood.

Cognition and memory require carefully balanced excitatory and inhibitory activity (Zhou and Yu, 2018). In different AD mouse models, impairments in inhibition precede plaque formation, disrupting brain rhythms associated with memory formation (Arroyo-García et al., 2021; Li et al., 2021; Nuriel et al., 2017; Sederberg et al., 2007). Modified inhibitory tone in early AD is likely related to changes in the intrinsic excitability of local circuit inhibitory interneurons. For example, AP firing is altered in ‘fast spiking’ PV interneurons in different human APP (hAPP)-expressing mice (Arroyo-García et al., 2021; Caccavano et al., 2020; Chen et al., 2018; Martinez-Losa et al., 2018; Petrache et al., 2019; Verret et al., 2012). Interestingly, altered PV physiology may occur before changes to other neighboring neuron subtypes (Hijazi et al., 2019; Park et al., 2020). Altered AP firing in PV cells could result from changes in the expression of genes that regulate excitability (Martinez-Losa et al., 2018). However, major shifts in gene and protein expression may only materialize after substantial plaque formation (Bundy et al., 2019) in AD. Thus, a systematic evaluation of molecular mechanisms contributing to altered firing in PV cells is required.

In this study, we used a viral-tagging method to examine PV interneuron excitability in the somatosensory cortex of young adult 5xFAD mice. PV interneurons from 5xFAD mice displayed strongly dampened firing near-threshold and modified action potential (AP) waveforms, indicating dysregulation of either Na^+^ or K^+^ channels. Combined examination of several AP firing parameters, computational modeling, and PV-specific qPCR indicated that changing Na^+^ channel availability was not responsible for changes in AP firing. However, we observed alterations in K^+^ channel activation and kinetics in AD mice, independent of changes in K^+^ gene expression. Using dynamic clamp and additional PV modeling, we found that these shifts in K^+^ channel activation could recapitulate the observed phenotypes in 5xFAD mice. Furthermore, K^+^ channel-induced changes in PV firing were sufficient to induce circuit hyperexcitability and modified gamma output in a reduced cortical model. Together, these results establish a causal relationship between ion channel regulation in PV interneurons and cortical circuit hyperexcitability in early AD, independent of changes in gene expression.

## Results

### Near-threshold suppression of AP firing in PV interneurons of young 5xFAD mice

To evaluate physiological phenotypes of PV interneurons in 5xFAD and wild type control mice, we implemented an AAV viral-enhancer strategy (Vormstein-Schneider et al., 2020) to specifically label PV interneurons. Mature animals were injected with this PV-specific vector (referred throughout as ‘AAV.E2.GFP’) in layer 5 somatosensory cortex before plaque formation (postnatal day 42-49) (Bundy et al., 2019; Jawhar et al., 2012; Li et al., 2021; Oakley et al., 2006). Acute slices were obtained ∼7 days later and GFP-expressing (GFP^+^) cells were targeted for patch clamp using combined differential contrast and epifluorescent imaging (Figure 1A). Current clamp recordings from wild type mice displayed high-frequency, non-adaptive repetitive spiking characteristics of PV cells (Figure 1B). In addition, the expression of several known PV interneuron genes was confirmed in AAV.E2.GFP^+^ neurons (Chow et al., 1999; Ogiwara et al., 2007; Rudy et al., 2011) using qPCR, the levels of which were indistinguishable from PV interneurons isolated in identical fashion from PV-Cre mice (Figure S1).

**Figure 1.**
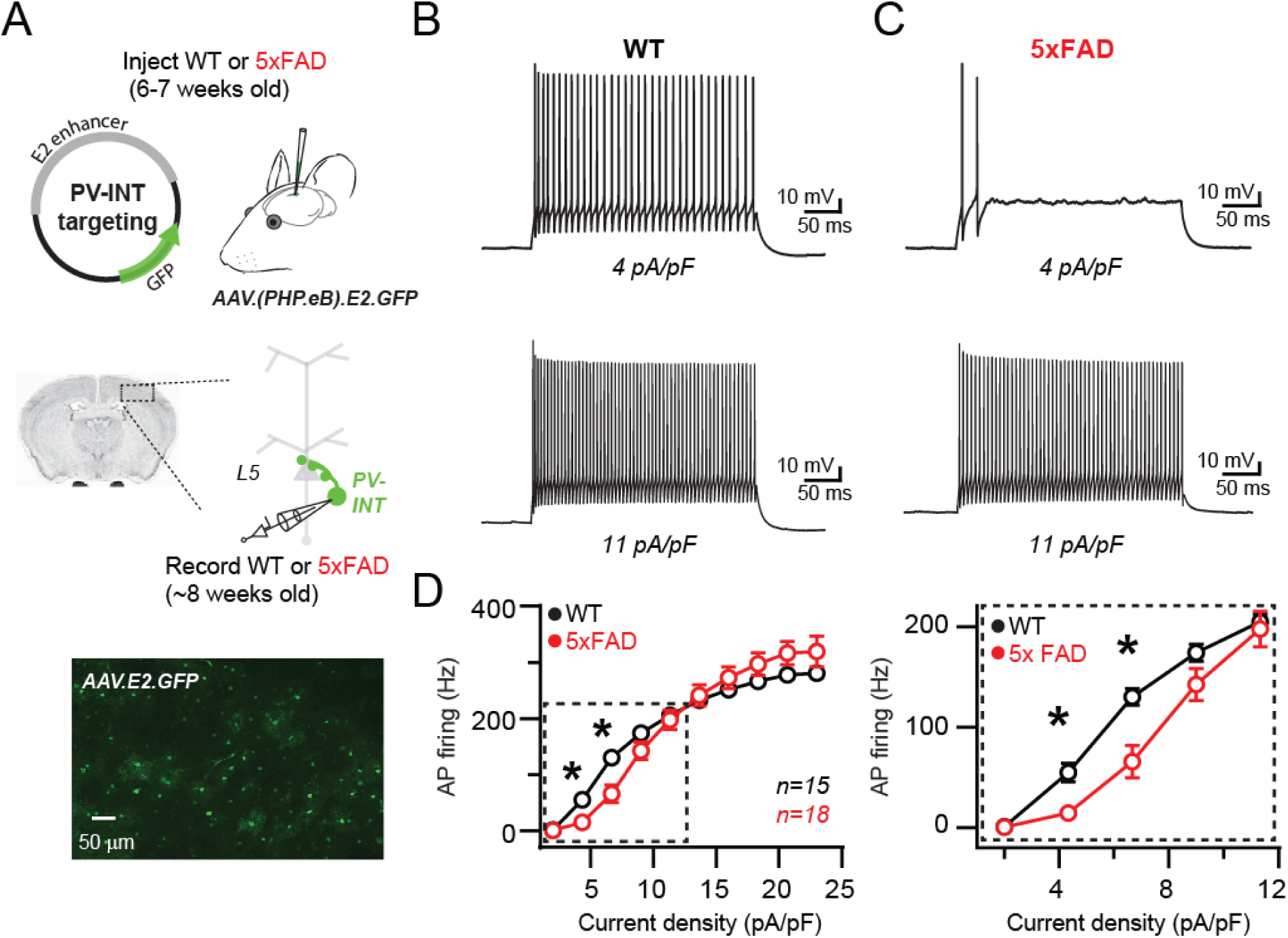
Reduced AP firing frequency in PV interneurons of young 5xFAD mice. (A) Graphical summary of AAV.E2.GFP stereotactic injection in somatosensory cortex and subsequent whole-cell current clamp recordings from GFP^+^ PV interneurons (PV-INT). (B) AP firing elicited in WT mice by square pulse current injections of varying magnitude normalized to cellular capacitance during recordings. (C) AP firing elicited in 5xFAD mice at current density levels matched to WT mice for comparison. (D) Group data summary of AP firing frequency in WT and 5xFAD mice. Significance was defined by RM two-way ANOVA (p < 0.05) with Sidak’s multiple comparison test). For all summary graphs, data are expressed as mean (± SEM). **See also** Figure S1.

Recent studies of several different hAPP expressing mouse models have demonstrated abnormal AP firing in GABAergic interneurons at different stages of plaque deposition (Hijazi et al., 2019; Mondragón-Rodríguez et al., 2018; Park et al., 2020; Petrache et al., 2019; Verret et al., 2012). In prodromal 5xFAD mice, we found that continuous spiking was severely dampened in layer 5 PV neurons in the near-threshold range, however, spike-frequency was unaltered near their maximal firing rate (Figure 1C & D). Passive parameters were unaltered when comparing WT and 5xFAD, including input resistance (94.9 ± 5.9 and 103.5 ± 8.4 MΩ; p = 0.83; unpaired t-test) and holding current immediately after break-in (17.5 ± 7.8 and 19.1 ± 10.5 pA; measured at −60 mV; p = 0.41; unpaired t-test), suggesting that an active mechanism was responsible for the observed differences in spike-frequency.

### Altered AP waveform and excitability are uncoupled from changes in Nav channels properties and mRNA expression

The extraordinarily rapid onset and repolarization of PV-APs depends on the combined expression of fast voltage-gated sodium (Na_v_) and potassium (K_v_) channel families (Baranauskas et al., 2003; Catterall et al., 2010; Cheah et al., 2012; Erisir et al., 1999; Goldberg et al., 2008; Gu et al., 2018; Rudy and McBain, 2001; Wang et al., 1998). Whether altered expression of voltage-gated channels emerge before plaque deposition is unclear. Changes in the expression of channels from the Na_v_1 family may contribute to altered spiking in cortical PV interneurons from hAPP-expressing FAD mice (Martinez-Losa et al., 2018; Verret et al., 2012; but see Saito et al., 2016). Therefore we examined parameters associated with fast-activating Na_v_ channels (Kole et al., 2008; Li et al., 2014; Platkiewicz and Brette, 2010) however, found no significant differences between 5xFAD and control mice (Figure 2A & B). AP afterhyperpolarization (AHP) amplitude was also unaltered (Figure 2B).

**Figure 2.**
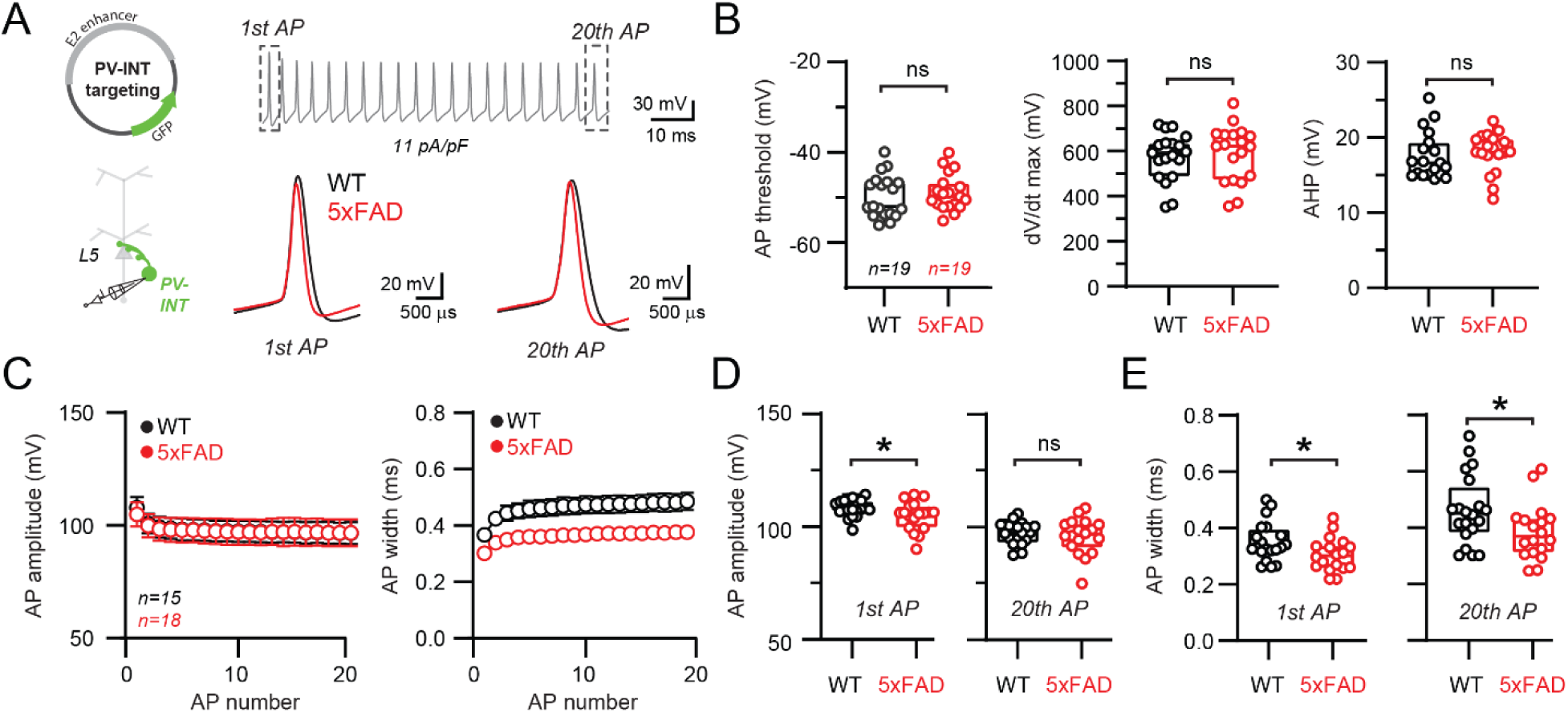
Altered AP waveforms in PV interneurons of 5xFAD mice. (A) AP waveforms and properties of GFP^+^ interneurons were compared at 11 pA/pF square pulse injections in WT and 5xFAD mice. In the enlarged view, APs from the 1^st^ and 20^th^ spike in the train of WT and 5xFAD mice are superimposed for comparison. (B) Summary data of AP properties. No differences in AP threshold, dV/dt maximum, or AHP were observed (p > 0.05; unpaired t-test). (C) Relationship between AP amplitude or width in WT and 5xFAD mice and AP # during spike trains elicited with 11 pA/pF current injection. Data are expressed as mean (± SEM). (D) Summary data of AP amplitude for the 1^st^ and 20^th^ APs in WT and 5xFAD mice. (E) Summary data of AP width for the 1^st^ and 20^th^ APs in WT and 5xFAD mice. For (B,D &E) individual data points and box plots are displayed. Significance was defined as p < 0.05; unpaired t-tests.

Na_v_ channel deficits result in reduced AP amplitude and contribute to AP failure during repetitive firing (Catterall et al., 2010; Escayg and Goldin, 2010; Gu et al., 2018; Wart and Matthews, 2006). Using a serendipitous current injection step where spike-frequency was indistinguishable between 5xFAD and control mice (11 pA/pF; Figure 2A), a subtle reduction in the amplitude of the initial AP was observed (Figure 2D). However, this reduction did not progressively worsen during continued firing (Figure 2C & 2D) as seen in mouse models where Na_v_1 channels were altered (Ogiwara et al., 2007; Yu et al., 2006). Interestingly, AP repolarization was more rapid across the entire spike train (quantified as a reduction in full AP width at half maximal amplitude; Figure 2C & E) in 5xFAD mice.

To test whether a Na_v_ channel mechanism could describe the AP firing phenotypes observed in 5xFAD mice, we built a simplified PV NEURON model constrained by our measurement AP parameters. Using the model, we independently simulated how changes in overall Na_v_ conductance, activation voltage, and kinetic properties affected relevant AP firing properties (Figure 3A). Significant reduction of Na_v_ conductance density (up to 50% of control) could lessen AP firing at near-threshold current steps (Figure 3B). However, this reduction was accompanied by complete firing failures at high frequencies (Verret et al., 2012) (Figure 3B), which was not observed in 5xFAD mice. Furthermore, AP width was unaltered over a broad range of Na_v_ conductance densities (Figure 3C) suggesting that AP width narrowing observed in 5xFAD mice was also due to a Na_v_-independent mechanism. In contrast, changing Na_v_ conductance density was associated with changes in AP threshold and maximal dV/dt (Figure 3D) which were unaltered in our recordings (Figure 2). Shifting Na_v_ kinetics or activation voltage also could not explain the observed 5xFAD phenotypes (Figure S2A-S2C).

**Figure 3.**
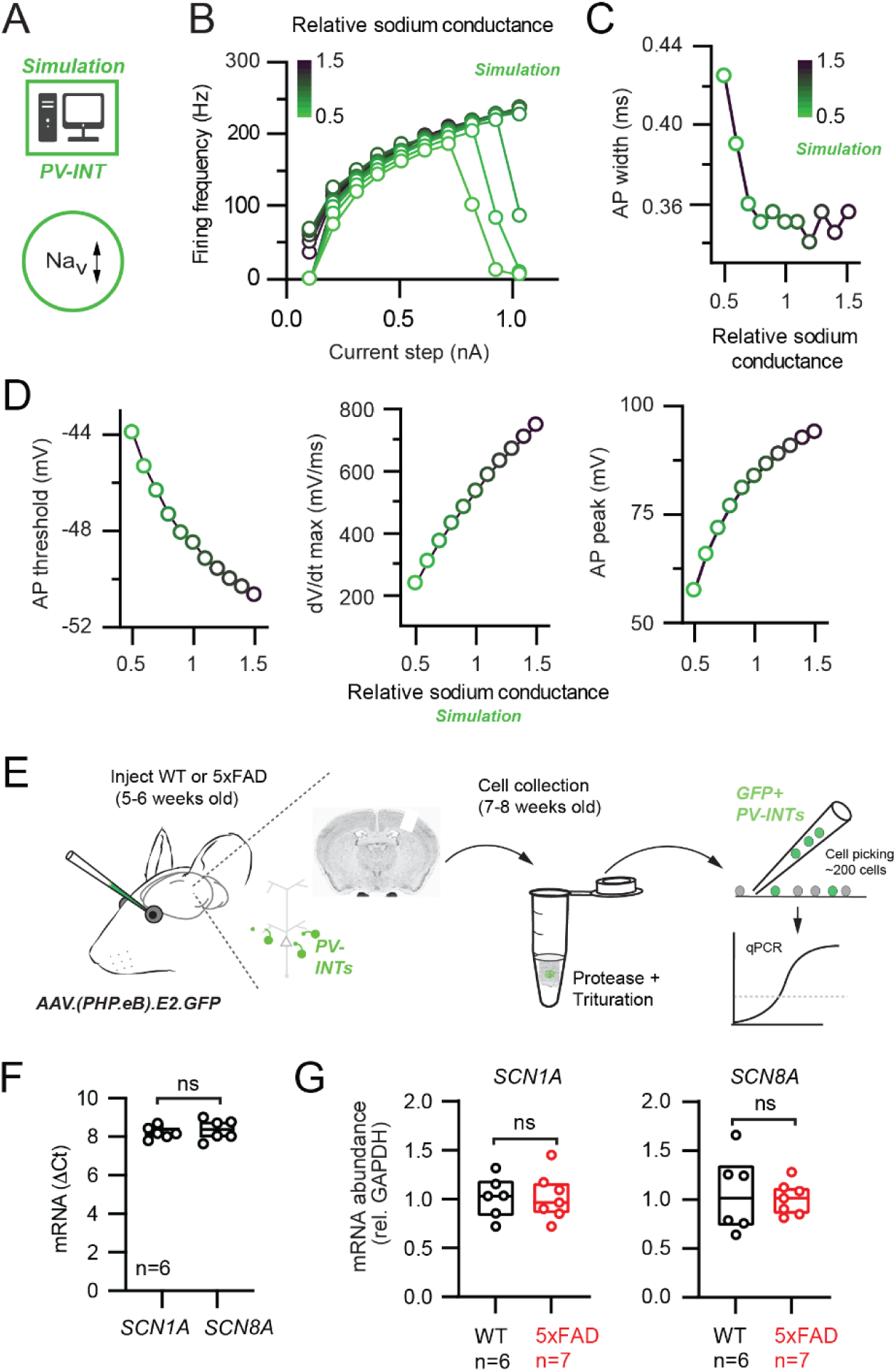
Na_v_ channel changes do not explain changes in PV interneuron excitability in 5xFAD mice. (A) Depiction of PV cell single compartmental model with modified Na_v_ channel properties. (B) Simulated relationship (S/cm^2^) between the magnitude of injected current and AP firing frequency at variable Na_v_ conductance densities. (C) Summary relationship of AP width and relative Na_v_ conductance density (± 50% from control Na_v_ conductance). (D) Summary graphs depicting the effect of changing Na_v_ conductance density on AP threshold, dV/dt maximum, and AP peak (± 50% from control Na_v_ conductance). (E) Depiction of cell-type-specific qPCR of *SCN1* genes following retro-orbital AAV injection in 4-6 week old mice. Individual neurons were physically isolated, hand-picked and pooled after allowing 2-3 weeks for cortical expression. (F) Comparative qPCR expression of *SCN1A* and *SCN8A* in WT mice. (G) Quantification of *SCN1A* and *SCN8A* mRNA expression between WT and 5xFAD mice. For (F) and (G) data are expressed as individual data points from each individual mouse with box plots superimposed. **See also** Figure S2.

To complement our Na_v_ modeling, we performed PV interneuron-specific qPCR (Tasic et al., 2018) by isolating and pooling AAV.E2.GFP^+^ neurons from dissected somatosensory cortex following AAV retro-orbital injection (Chan et al., 2017) in 5xFAD and control mice (Figure 3E). Expression of Na_v_1.1 (*SCN1A*) and Na_v_1.6 (*SCN8A*) were detected in wild type PV interneurons (Figure 3F). Relative to control, no changes in mRNA expression of either subunit in 5xFAD mice was found. (Figure 3G). Together, our patch clamp recordings, simulations, and gene expression data indicate that modifications in Na_v_ channel expression cannot account for the observed changes in PV firing in our pre-plaque hAPP model.

### Biophysical but not gene expression changes of Kv3 channels in PV interneurons

The distinct firing phenotype and rapid AP repolarization of fast-spiking PV cells require expression of fast-activating K_v_ channels, which complement Na_v_1 (Gu et al., 2018). Thus, by ruling out Na_v_ channels as viable candidates for explaining the above differences, we postulated that altered K_v_ channel availability could contribute to AP firing differences observed in 5xFAD mice. K_v_3 channels are highly expressed in PV cells, and possess extremely fast kinetics which set AP width and firing rate in fast-spiking neurons (Barry et al., 2013; Erisir et al., 1999; Rowan et al., 2014; Song et al., 2005). To record K_v_3 conductances from PV interneurons, we obtained outside-out patches from AAV.E2.GFP^+^ neurons in both 5xFAD and control mice. Patches were puffed with TEA (1mM) to block and *post-hoc* isolate K_v_3 currents (Figure 4A). Certain calcium dependent potassium channels (i.e., B_K_) may also be sensitive to 1mM TEA (Coetzee et al., 1999); however, fast-spiking interneurons do not utilize B_K_ to shape APs (Casale et al., 2015; Rowan et al., 2014). Large TEA-sensitive currents were isolated in patches from PV cells (Figure 4B) displaying characteristic K_v_3-like properties including a relatively depolarized half-activation voltage (Figure 4D) and submillisecond activation kinetics (Figure 4E) (Baranauskas et al., 2003; Lien et al., 2002; Rudy and McBain, 2001).

**Figure 4.**
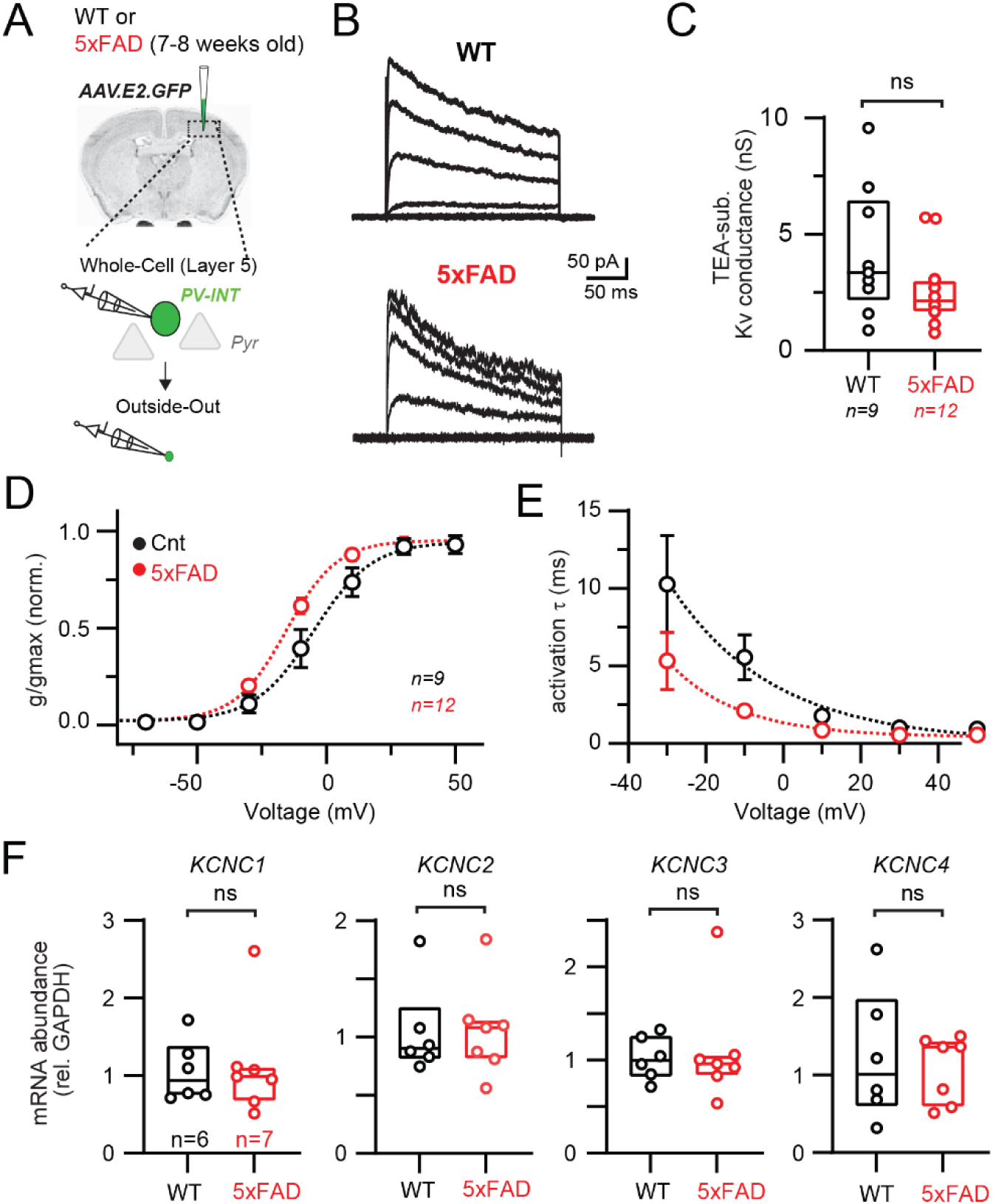
Modified K_v_3 channel biophysics in 5xFAD mice. (A) Experimental workflow for obtaining outside out patches from PV interneurons in WT and 5xFAD mice. (B) Representative K_v_3 currents isolated from outside out patches in WT and 5xFAD mice. (C) Data summary of maximal K_v_3 conductance in WT and 5xFAD mice (p > 0.05; unpaired t-test). Individual data points from each patch and box and whisker plot summaries are displayed. (D) Summary of activation voltage of K_v_3 conductance isolated from patches in WT and 5xFAD mice. Conductance was normalized to the maximal overall conductance (gmax) for each cell. The average dataset was fit with a Boltzmann function with individual values expressed as mean (± SEM). (E) Summary of activation time constant (τ) of K_v_3 currents in isolated from patches in WT and 5xFAD mice. Datasets were fit with single monoexponential decay functions and are expressed as mean (± SEM). (F) Comparison summary of *KCNC*1-4 mRNA expression between WT and 5xFAD mice from isolated and pooled PV interneurons. Individual data points from each mouse and box plot summaries are displayed. No differences were found between WT and 5xFAD cohorts for any of the four subunits (p > 0.05; unpaired t-tests).

Substantial changes in K_v_3 channel availability may account for the observed differences in AP firing in 5xFAD mice (Figure 1). However, the overall TEA-subtracted conductance was unchanged in 5xFAD (Figure 4C). The proportion of TEA-insensitive K_v_ conductance was also unchanged (wild type, 33.1 ± 2.9%; 5xFAD, 33.0 ± 2.3%; p = 0.98; unpaired t-test; n = 9 and 12; respectively). Interestingly, we observed differences in the biophysical properties of K_v_3 channels in the 5xFAD model. Channels were found to activate at more hyperpolarized (left-shifted) voltages (Figure 4D; half activation voltage −6.6 mV control vs −15.5 mV in 5xFAD). Furthermore, K_v_3 activation tau increased across the observable range in 5xFAD mice (Figure 4E). Moderate K_v_3 inactivation was observed (Alle et al., 2011) in both control and 5xFAD mice (peak to steady state amplitude ratio (Hu et al., 2010); 0.59 ± 0.07 and 0.53 ± 0.04, control and 5xFAD respectively; p = 0.43; unpaired t-test; n = 9 and 12; respectively).

Differential expression of the four known Kv3 channel *KCNC* subunits in 5xFAD mice could account for the observed shifts in K_v_3 biophysics (Figure 4 D & E). To evaluate this possibility, we again performed PV interneuron-specific qPCR by isolating AAV.E2.GFP^+^ cells (Figure 4F) as described earlier. Expression of all four subunits was confirmed in PV cells from somatosensory cortex, however, no differences in mRNA expression were found between 5xFAD and control mice, for any of the four *KCNC* subunits (Figure 4F). Together these data indicate that the modifications responsible for divergent biophysical properties of K_v_3 channels occur via post-translational regulation at this pre-plaque disease stage.

### Modified K_v_3 channel biophysics recapitulate the 5xFAD phenotypes in a PV model

To test whether modifying K_v_3 channel biophysics alone could adequately explain the AP firing phenotypes in 5xFAD mice, we returned to our reduced PV cell simulation (Figure 5A). In control conditions, our model PV neuron increased firing in relation to the magnitude of current injection (Figure 5B & 5C). Notably, when the K_v_3 membrane potential dependence was hyperpolarized as observed in 5xFAD PV neurons (Figure 5A & 5B; control absolute half activation voltage = 5.0 mV; absolute test Vshift (−10 mV) = −15.0 mV) we found that AP ring was strongly dampened in he near-threshold range (Figure 5B & 5C; see also Lien and Jonas 2003), mirroring changes in 5xFAD mice. Shifting K_v_3 activation voltage left-ward also led to slight reduction in firing frequency at higher current injection levels, which could be normalized with a concurrent K_v_3 increase in the activation tau (Figure 5C). Modulation of K_v_3 kinetics alone could also modify maximal firing frequency in either direction (Figure 5D).

**Figure 5.**
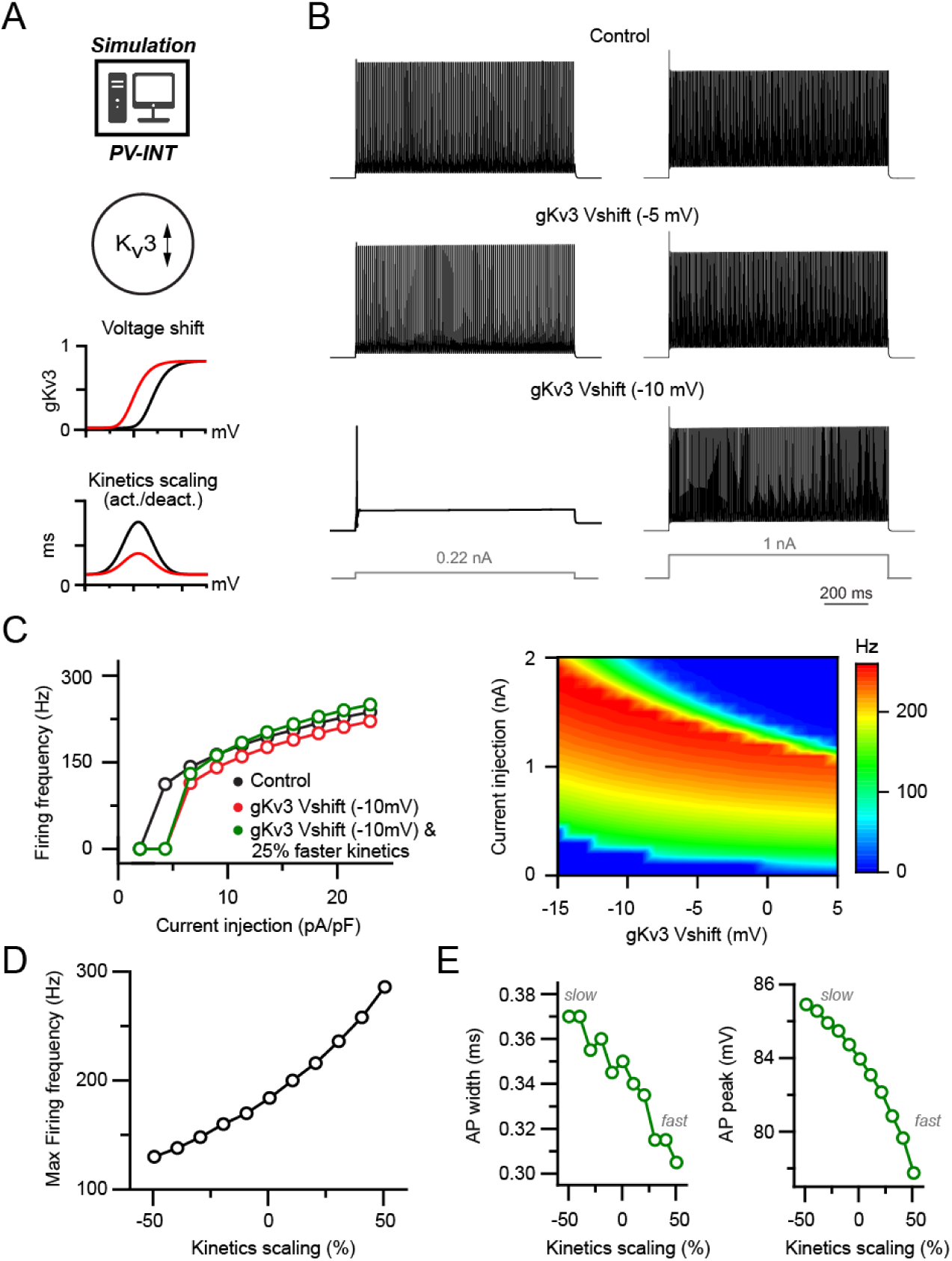
Effect of biophysical K_v_3 dysregulation on AP firing in a PV model. (A) PV cell single compartmental model with modified K_v_3 channel properties. K_v_3 activation voltage and kinetics were independently or simultaneously modified in the following simulations. When applied, activation and deactivation kinetics were scaled together (± 50% of control). (B) AP firing elicited by square pulse current injections at control and hyperpolarized K_v_3 activation voltages. Two example current injection magnitudes are displayed. (C) Summary data of firing frequency changes in different simulated K_v_3 conditions. Near-threshold AP firing is reduced with hyperpolarized K_v_3 activation independent of increased K_v_3 kinetics. (D) Effect of modifying K_v_3 channel kinetics (± 50% of control) alone on maximal firing frequency in PV neuron compartmental model. (E) Effect on K_v_3 channel kinetics changes on simulated AP width and amplitude.

AP repolarization is differentially shaped by distinct kinetic properties of different K_v_ subtypes (Bean, 2007; Dodson et al., 2002; Pathak et al., 2016; Rowan et al., 2014; Wang et al., 1998). As AP width in our PV cell model was largely uncoupled from changes in Na_v_ conductance, we hypothesized that AP width was also influenced by changes in K_v_3 channel kinetics (Baranauskas et al., 2003). Indeed, increased activation kinetics (tau) were correlated with a reduction in AP width which could also influence AP amplitude (Figure 5E). Hence upon model exploration of the relevant K_v_3 biophysical parameters, we could fully recapitulate the AP firing phenotypes observed in 5xFAD PV cells.

### Introduction of modified K_v_3 conductance reproduces near-threshold hypoexcitabilty in PV interneurons

While powerful, our PV model predications are based on simplified biophysical information. To increase confidence that altered K_v_3 channel properties can explain reduced near-threshold excitability in intact PV neurons, we employed an Arduino-based dynamic clamp system (Desai et al. 2017). Dynamic clamp allows for real-time injection of current constrained by predefined voltage-gated conductances, such as gK_v_3, during current clamp recordings (Figure 6A). Furthermore, distinct properties (e.g., activation voltage) of these conductances can be adjusted online during recordings.

**Figure 6.**
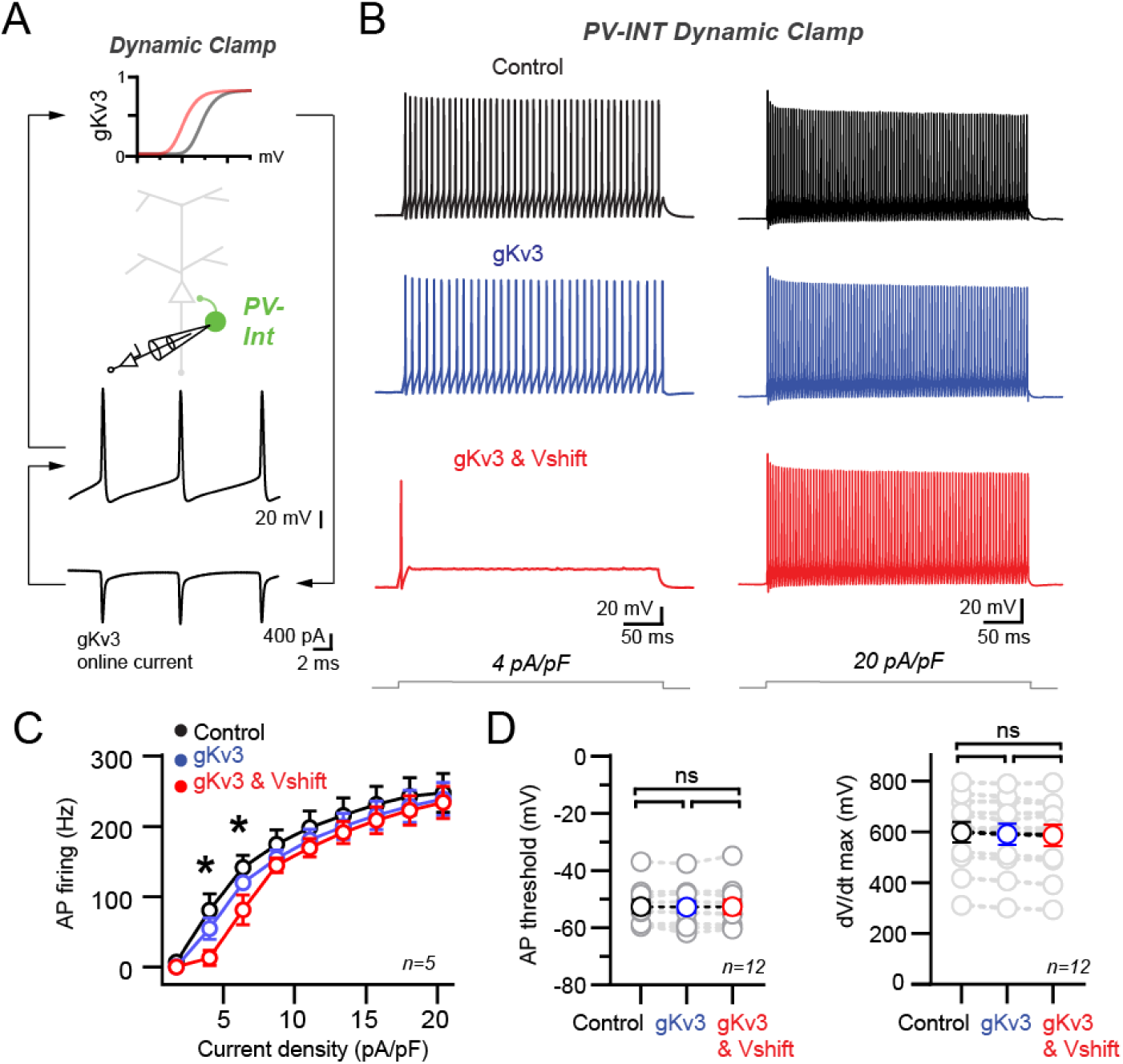
Recapitulation of the 5xFAD phenotype in PV cells using dynamic clamp. (A) Targeted dynamic clamp recordings from an AAV.E2.GFP_+_ neuron. Online K_v_3 response (20 nS online gKv3) shown during AP firing in a PV interneuron. (B) AP firing responses to two different square pulse current injection levels in three distinct K_v_3 dynamic clamp conditions in the same cell. (C) Summary data plot across a range of current injections from dynamic clamp conditions. Statistical significance was tested between the gKv3 (blue) and gKv3 & Vshift (red) conditions by RM two-way ANOVA (p < 0.05) with Sidak’s multiple comparison test. (D) Summary plots for AP threshold and dV/dt maximum in each of the dynamic clamp conditions tested within each cell. No differences were observed in any condition using RM one-way ANOVA (p < 0.05) with Tukey’s multiple comparison test. In all datasets individual values are expressed as mean (± SEM).

Dynamic clamp recordings were performed in targeted recordings from AAV.E2.GFP^+^ neurons in wild type mice. AP firing was examined in three distinct K_v_3 conditions (Figure 6B; *Control* [no dynamic clamp conductance added, 0.0 nS *gKv3*]; *+gK_v_3* [absolute half-activation voltage, −5.0 mV]; and *+gK_v_3 & Vshift* [‘5xFAD’ absolute half-activation voltage, −15.0 mV]). Modest supplementation of additional wild type K_v_3 conductance (*+gK_v_3_;_* 20 nS) had no discernable effect on AP firing across a range of current densities (Figure 6B & 6C). However, introduction of an identical magnitude of the 5xFAD-modeled K_v_3 conductance (*+gK_v_3 & Vshift*; 20 nS) induced a specific reduction in near-threshold firing without affecting high-end frequencies (Figure 6B & 6C). Compared to control, AP threshold and dV/dt maximum were unchanged in both *gKv3* test conditions (Figure 6D). Together with our NEURON simulation data, these dynamic clamp recordings indicate that introduction of biophysically modified K_v_3 conductance can reproduce the hypoexcitable phenotype observed in PV interneurons in prodromal 5xFAD mice.

### K_v_3 modulation reduces synaptically-evoked AP firing in PV interneurons

*In vivo*, cortical PV neurons often fire at the lower end of their dynamic range (Yao et al., 2020; Yu et al., 2019). To examine how K_v_3 channel modulation affects PV interneuron firing in a realistic network condition, we imposed several hundred sparsely active (see Methods) excitatory and inhibitory synapses onto our PV NEURON simulation (Figure 7A). In control conditions, the PV cell fired regularly (30.64 ± 0.39 Hz). Hyperpolarization of the control K_v_3 membrane potential dependence was inversely correlated with spike-frequency (Figure 7B).

**Figure 7.**
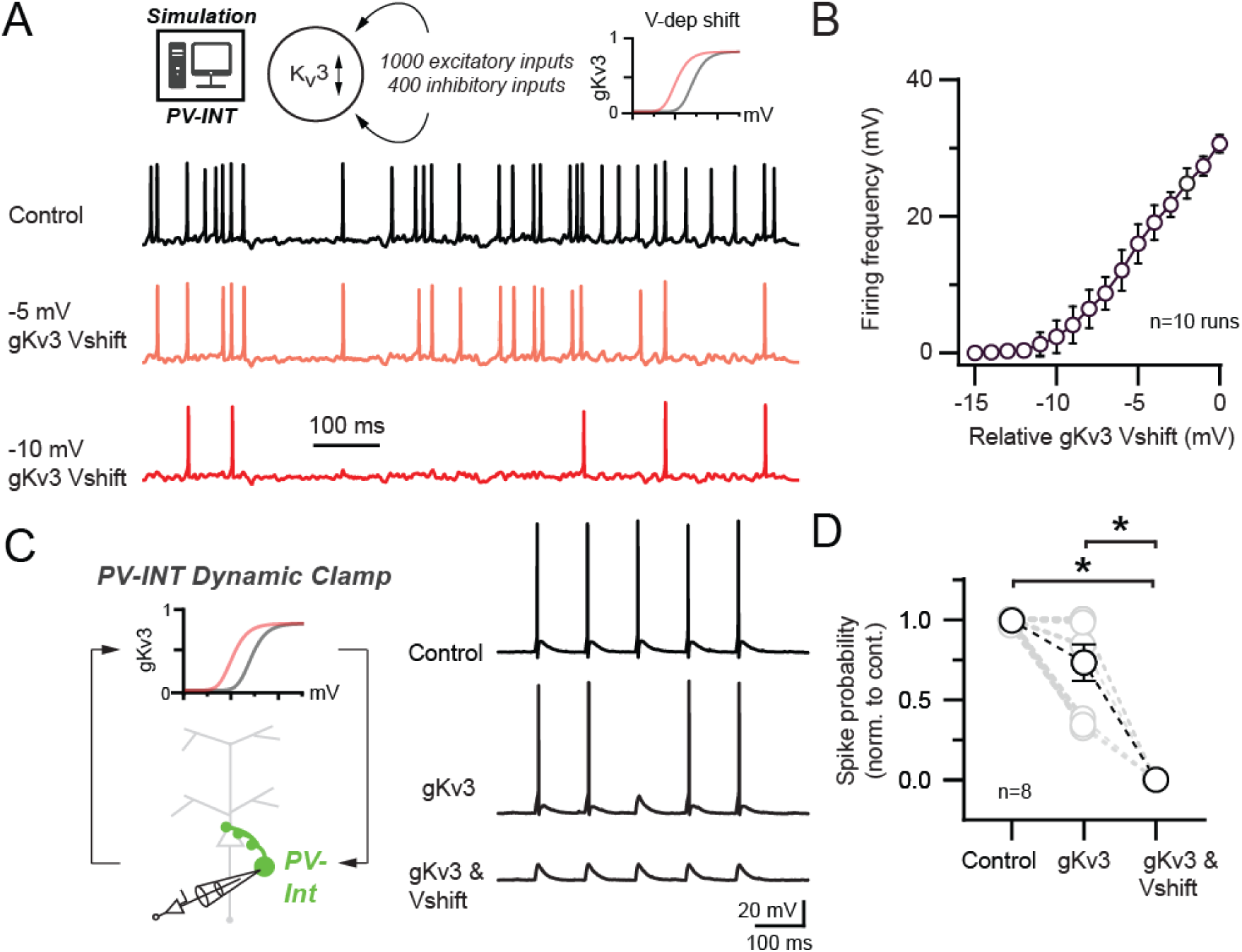
Effect of 5xFAD-related K_v_3 channel modulation on synaptically evoked AP firing. (A) Simulated responses of PV cell compartmental model with continuous excitatory and inhibitory inputs in control and with hyperpolarized K_v_3 activation voltages. (B) Summary graph of PV compartmental model firing frequency in response to continuous synaptic inputs at increasingly hyperpolarized K_v_3 activation voltages. 0 mV represents the relative control K_v_3 activation voltage. (C) 10 Hz gEPSP-evoked AP firing in dynamic clamp recordings from AAV.E2.GFP^+^ neurons in acute slice. In control conditions, gEPSP conductance was calibrated such that the majority of stimuli evoked APs. Within recordings the gEPSP amplitude was constant while the cell was subjected to varying gK_v_3 dynamic clamp conditions. (D) Spike probability summary in response to gEPSPs in varying gK_v_3 dynamic clamp. Significance was defined by one-way ANOVA (p < 0.05) with Tukey’s multiple comparison test). For all summary graphs, data are expressed as mean (± SEM). **See also** Figure S3.

Using dynamic clamp in wild type mice, we next sought to understand whether K_v_3 channel regulation could also diminish synaptically-evoked AP firing in intact PV (AAV.E2.GFP^+^) interneurons. *In vivo*, single excitatory synaptic inputs can reliably drive AP firing in PV neurons (Jouhanneau et al., 2018). Thus, we injected PV neurons with an excitatory conductance (gEPSP) (Sharp Marder 1993, Jaeger 2011, Xu-Friedman Regehr 2005) to reliably evoke AP firing at 10 Hz (gEPSP, 4.7 ± 1.0 nS; Figure 7C). Dynamic clamp addition of wild type K_v_3 conductance (*+gK_v_3*; 20 nS) had a non-significant effect on gEPSP-evoked AP firing (Figure 7C & D). Interestingly, injection of the 5xFAD-modeled K_v_3 conductance (*+gK_v_3 & Vshift*; 20 nS) strongly reduced gEPSP-evoked firing (Figure 7C and 7D).

While often referred to as high-voltage activating channels, K_v_3 channels open in the subthreshold range in GABAergic interneurons (Rowan and Christie, 2017) and regulate the magnitude of EPSPs (Hu et al., 2010). In PV NEURON simulations, hyperpolarizing the K_v_3 activation voltage could efficiently reduce the amplitude of EPSPs (Figure S3A), thus necessitating an increase in excitatory synaptic conductance to evoke an AP (Figure S3B). This modulation was also observed in further dynamic clamp PV recordings with subthreshold gEPSPs (3.6 ± 0.8 nS; Figure S3C). Together, these data argue that enhanced subthreshold activation of K_v_3 contributes to near-threshold PV hypoexcitabilty during early-stage AD.

### Modulation of PV K_v_3 channels elicits network hyperexcitability in a reduced layer 5 circuit model

Precisely timed synaptic inhibition of neuronal circuits provided by PV interneurons is indispensable for network operations (Cardin et al., 2009; Da et al., 2012; Fuchs et al., 2007; Sohal et al., 2009). In order to understand the network consequences of the observed PV phenotype in young 5xFAD mice, we developed a local PV-PC network model (Figure 8A). Connection strengths and probabilities for the network consisting of 200 PCs and 20 PV cells were based on previous reports (Bock et al., 2011; Galarreta and Hestrin, 2002; Hofer et al., 2011; Markram et al., 2015; Perin et al., 2011). The model reproduced key features of local PV circuit models including gap-junction related firing synchrony (Wang and Buzsáki, 1996) and recurrent connection related synchrony (Bartos et al., 2007).

**Figure 8.**
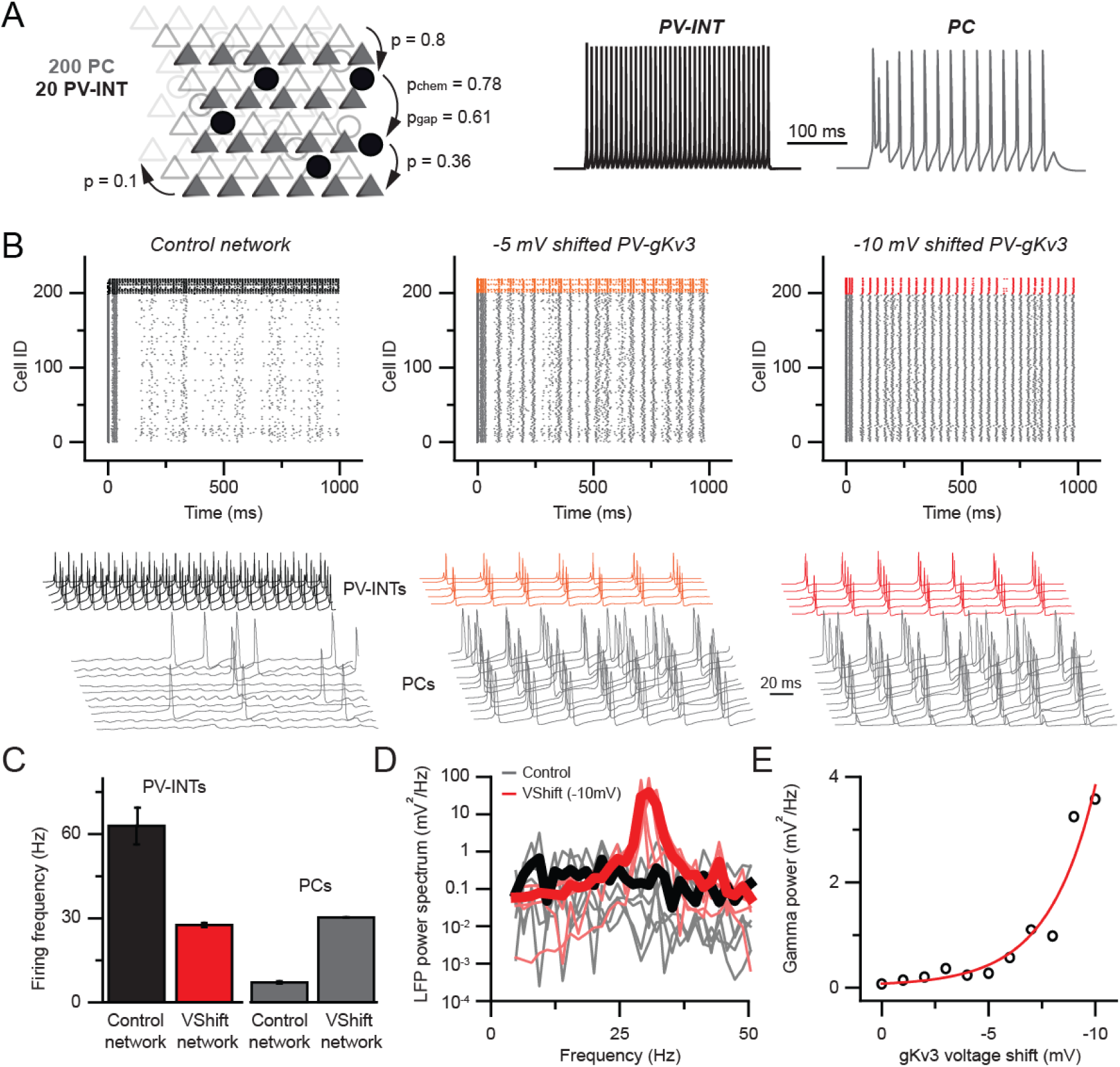
Hyperexcitability and increased gamma following PV-specific Kv3 modulation. (A) Simplified cortical network consisting of 200 pyramidal cells (PC; triangles) and 20 PV (circle) cells. Connection probabilities between and within cell groups are set based on literature. 300 ms long spiking responses for single PC and PV cells are shown on the right. (B) Raster plots depicting 1 s long network activity of the 220 cells in the network. The top 20 cells correspond to PV cells (black, orange, red), bottom 200 cells show PC activity (grey). The effect of relative −5.0 and −10.0 mV shifts in gK_v_3 of PV cells are compared to control. Representative traces are shown from 5 PV cells and 10 PC. (C) Mean firing frequency of PV cells and PCs upon −10 mV relative voltage shift of gK_v_3 in PV cells. Data are expressed as mean (± SEM). (D) Calculated local field potential (LFP) between 5 and 50 Hz, produced by 220 cells in the network. The activity level of individual cells was randomized and network simulations were repeated 5 times in control conditions and with a −10 mV relative shift in gK_v_3 of PV cells. Individual LFP traces are shown in light grey and light red. Mean LFP traces are shown in bold black and red. (E) Gamma power in relation to the voltage shift of gK_v_3 in PV cells. Gamma power was calculated by averaging LFP signals between 30 and 50 Hz. Continuous red line depicts the exponential relationship between the two variables. **See also** Figure S4.

We found that gradual shifting of the voltage dependence of gK_v_3 conductance in PV cells markedly increased the firing rate of the simulated PCs (Figure 8B, control: 7.07 ± 0.42 Hz, 10 mV; Vshift: 30.3 ± 0.12 Hz, n = 200, p < 0.0001, paired t-test). This network hyperexcitability can be attributed to the altered excitation-inhibition ratio due to the effects of gKv3 biophysical changes of PV interneuron firing. Specifically, in the control network, PV firing (62.9 ± 6.58 Hz mean firing, n = 20) was constrained by their recurrent connections, gap junctions and sporadic entrainment by the PC population’s low firing rate. However, when the excitability of PV cells was dampened by altered gK_v_3 voltage-dependence (Figure 8C; n = 20 runs), PCs were released from the high inhibitory tone resulting in network hyperexcitability, which is a hallmark of recurrently connected pyramidal cells networks (Morgan Soltesz 2007, Paz Huguenard 2015).

Next, we investigated whether the increase in network excitability resulted in altered oscillatory behavior. We found that there was a significant increase of gamma power at 30 Hz (Figure 8D, 0.13 ± 0.08 and 38.7 ± 14.76 mV^2^/Hz, n = 5 each, p < 0.05, paired t-test; for control and shifted gK_v_3 network respectively) which is in agreement with previous work (Sohal et al., 2009).

Our simulations demonstrate that alterations in the voltage dependence of a single PV conductance can have substantial effects on local network activity. However, minor deviations from the ensemble mean can arise from the stochastic nature of channel opening and closing (Cannon et al., 2010; Lemay et al., 2011) and from interactions with auxiliary channel subunits (Oláh et al., 2020; Yu et al., 2005). Therefore, we tested the stability of the network upon perturbations of gK_v_3 gating. Our results showed an exponential relationship (R^2^ = 0.93) between the voltage shift of gK_v_3 in PV cells (Figure 8E) and network gamma power. This nonlinearity indicates that although a ∼10 mV shift can alter circuit behavior, the network is protected against expected stochastic ion channel fluctuation-induced alterations in excitability. Together, our results demonstrate that a hypersynchronous (Figure S4) and hyperactive network activity can emerge as a consequence of altered PV interneuron K_v_3 biophysics.

## Discussion

In this study, we report a novel mechanism contributing to cortical circuit dysfunction in an early-stage AD mouse model. Our findings indicate that posttranslational modulation of K^+^ channel biophysics contributes to cortical PV interneuron dysfunction in early AD. In a simplified circuit model, this K^+^ channel mechanism caused cortical network hyperexcitability and modified signaling specifically in the gamma frequency domain. Our results represent a novel cellular mechanism with a causal link to overall circuit hyperexcitability, thus presenting a potential therapeutic avenue to combat AD progression in its early stages.

### PV interneuron pathophysiology in AD models

PV-positive GABAergic interneurons constitute a substantial proportion (∼ 40%) of the total cortical interneuron population (Tremblay et al., 2016). These interneurons form powerful inhibitory synapses with local pyramidal neurons, thereby regulating a variety of cognitive functions (Yao et al., 2020). In several different AD mouse models, investigators have observed abnormal PV intrinsic excitability, however, mechanistic understanding of this phenomenon is incomplete. Here we report reduced cortical PV firing in the 5xFAD model. In complementary AD mouse models, human APP and PS1 proteins (e.g., APP/PS1, hAPPJ20) are also expressed at high levels and include mutations resulting in increased amyloid production. Within these models, PV interneurons display physiological phenotypes including altered AP firing (Hijazi et al., 2019; Verret et al., 2012). Notably, PV neurons were found to be more susceptible to shifts in their excitability with respect to neighboring pyramidal neurons in these studies. PV-specific vulnerability could manifest as a result of their high metabolic demand (Ruden et al., 2021) or through abnormal regulation of ion channel subunits necessary for maintaining their fast-spiking nature (Martinez-Losa et al., 2018).

Related changes in PV neuron excitability are evident among the hAPP mouse models. In layer 5 PV cells, we observed reductions in near-threshold AP firing and AP width, but AP amplitude and passive properties were largely unaffected. In hippocampal CA1 from 5xFAD mice, AP firing during synaptic recruitment was also strongly reduced (Caccavano et al., 2020). In layer 2/3 PV neurons of hAPPJ20 mice, overall AP firing rates were unchanged but a significant reduction in AP amplitude was observed (Verret et al., 2012), however, in hAPPJ20 hippocampal CA1, spike frequency was strongly reduced (Mondragón-Rodríguez et al., 2018). A CA1 study from APP/PS1 mice observed reduction in AP width but increased AP frequency (Hijazi et al., 2019). In next-generation hAPP KI mice, which express the hAPP at far lower levels with respect to the aforementioned APP models, PV firing frequency was also reduced in entorhinal cortex before plaque deposition (Petrache et al., 2019). Variations among these studies could depend on the disease severity at which observations were made, regional differences, or genetic differences between models. Nonetheless the related phenomena evident across these studies suggests that a unifying set of molecular mechanisms may spark circuit-level dysfunction in early AD.

### Mechanisms of altered PV excitability in AD

In a hallmark set of studies, differential expression of voltage gated Na^+^ channels in PV neurons was linked with network hyperexcitability in hAPP-expressing AD mice (Martinez-Losa et al., 2018; Verret et al., 2012). It is unclear whether other channel types are regulated and contribute to PV neuron dysfunction in AD. In this study we observed physiological changes in 7-8 week old 5x FAD mice, however, no detectable proteomic changes are expected up to 4 months of age in this model (Bundy et al., 2019). In keeping with this finding, we did not observe differences in Na_v_1 or K_v_3 mRNA levels in 7-8 week old mice. Rather, our results indicate that biophysical modulation of K_v_3 channels was responsible for reduced AP firing and AP width. Interestingly, reduced AP width was observed in PV cells before other intrinsic alterations in APP/PS1 mice (Hijazi et al., 2019) suggesting that K_v_3 modulation could precede that of other channels or homeostatic responses. Additional longitudinal studies at multiple stages of the disease will be necessary to parse out the emergence of cell-type-specific mechanisms during the disease.

Several plausible AD-related cellular processes could explain the biophysical modulation of K_v_3 observed in this study. Increases in β-Secretase (BACE1) activity in AD (Roßner et al., 2006), which cleaves full-length APP, promotes surface expression of K_v_3.4 subunits. K_v_3.4 subunits activate more rapidly and display hyperpolarized activation voltage in interneurons (Baranauskas et al., 2003; Rowan et al., 2016) similar to the K_v_3 phenotype observed here in 5xFAD mice. Alternatively, BACE1 could modulate the gating properties of membrane bound Kv3 channels (Hessler et al., 2015). A intermediate transmembrane protein product produced following BACE1 cleavage, called C99, can also regulate K_v_ channel activity (Manville and Abbott, 2021). Finally, increased levels of extracellular Aβ may regulate K_v_ channel conductance through direct interactions, or indirectly via channel phosphorylation (Farley et al., 2021). One or more of these hAPP-related processes could contribute to the K_v_3 channel dysregulation observed in 5xFAD mice here. Implementing the PV-type-specific viral approach utilized in this study in various AD models will allow for a deeper evaluation of these possibilities *in vivo* in future work.

### Relationship of PV interneuron dysfunction and circuit level disruptions

Circuit hyperexcitability is a prodromal indicator in familial and late-onset AD (Dickerson et al., 2005; Hämäläinen et al., 2007; Miller et al., 2008; Sperling et al., 2010; Busche and Konnerth, 2015; Lamoureux et al., 2021; Minkeviciene et al., 2009; Nuriel et al., 2017; Quiroz et al., 2010; Sepulveda-Falla et al., 2012). Altered PV interneuron firing occurs at early stages of the disease (Hijazi et al., 2019; Petrache et al., 2019), likely contributing to epileptiform activity and overall circuit hyper-synchrony in cortex. Using a layer 5 cortical circuit model, we found that PV-specific K_v_3 channel dysfunction resulted in overall hyperexcitability (Busche 2008; Palop and Mucke 2010).

Several PV cell-specific cellular and connectivity features, such as short input integration time-window (Hu Jonas 2014, Geiger Jonas 1997), frequent recurrent connections, and extensive gap junction coupling (Galarreta and Hestrin 2002) help regulate cortical circuit operations. PV cells are particularly important for maintaining signaling in the gamma frequency domain (Bartos et al., 2007). In our 5xFAD simulation, which produced near-threshold reduction in PV firing, we observed a sharp increase in gamma power that scaled with the severity of K_v_3 modulation. Similarly, reduced PV excitability can amplify gamma power in different cortical areas (Picard et al., 2019) likely through disruption of feedback inhibitory circuits (Sohal et al., 2009). Notably, increased gamma power was observed in AD patients during resting states (Wang et al., 2017). In the context of these studies, it is temping to hypothesize that near-threshold changes in PV firing may disrupt feedback inhibitory circuits in cortex in times of sparse coding. Conversely, reduction of PV excitability can also result in reduced gamma power in different contexts (Carlén et al., 2012). Thus bidirectional, PV-specific modulation of the gamma range is likely to be circuit and context dependent (Sohal and Verret 2012). The tendency for local gamma power to increase or decrease in different circuits in AD should provide insight into PV-specific cellular pathology.

Further disentanglement of the mechanisms of interneuron dysfunction in distinct AD models is necessary. Specifically, the relationship of hAPP, amyloid (Johnson et al., 2020; Rodgers et al., 2012), and its intermediate products to PV-related dysfunction and abnormal circuit function. The versatility and efficiency provided by the cell-type-specific enhancer approach used here can be implemented in future studies on novel AD mouse models, or by transgene expression through viral delivery (Kim et al., 2013), as well as in iPSC derived human neurons.

## Author Contributions

Conceptualization, M.J.M.R; Methodology, M.J.M.R., J.D., V.J.O and A.M.G.; Investigation, M.J.M.R., V.J.O and A.M.G; Writing – Original Draft, M.J.M.R. and V.J.O.; Funding Acquisition, M.J.M.R. and J.D; Supervision, M.J.M.R.

## Acknowledgments

We sincerely thank Niraj Desai and Dan Johnston (UT Austin) for their help in constructing the dynamic clamp system. A subset of AAV viral vectors were packaged in the Emory Viral Vector Core Facility. cDNA extraction and qPCR were performed with support from the Emory Integrated Genomics Core Facility. This work was supported by NIH grants R56-AG072473 (M.J.M.R.), R01-MH111529 (JD), and UG3MH120096 (JD); The Emory Alzheimer’s Disease Research Center Grant 00100569 (M.J.M.R.); Autifony Therapeutics (M.J.M.R); Simons Foundation Award 566615 (JD).

## Declaration of interests

The authors have no conflicts of interest to declare.

## Supplementary Information

**Figure S1, related to Figure 1.**
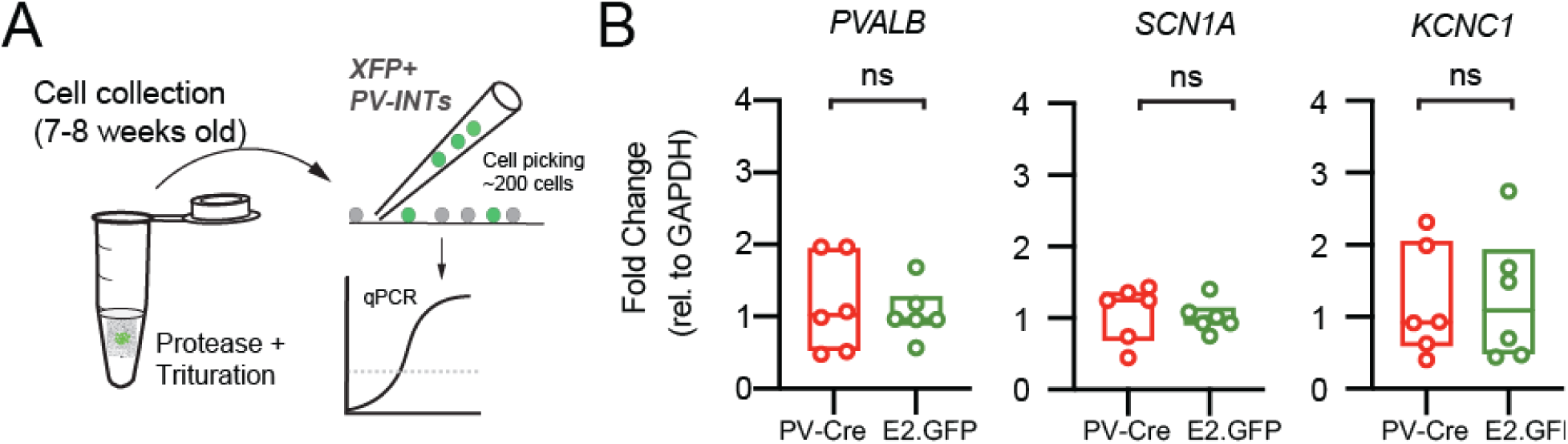
Confirmation of PV interneuron gene expression in AAV.E2.GFP^+^ neurons. (A) Depiction of cell-type-specific qPCR following stereotactic AAV injection in 5-6 week old mice. ∼200 GFP^+^ neurons were physically isolated and hand-picked from the somatosensory cortex at 7-8 weeks of age. (B) PV-specific gene expression was compared in two mouse strains. One cohort of PV-Cre mice (n=6) acting as the control group were injected with an AAV expressing a floxed tdTomato construct. A second test cohort (n=6) of WT mice were injected with AAV.E2.GFP. Expression of three known cortical PV interneuron-specific genes (*PVLAB*, *SCN1A*, *KCNC1*) were quantified for each cohort. There were no differences (p > 0.05; unpaired t-tests) between the two groups for any of these genes. Individual data points and box plots are displayed.

**Figure S2, related to Figure 3.**
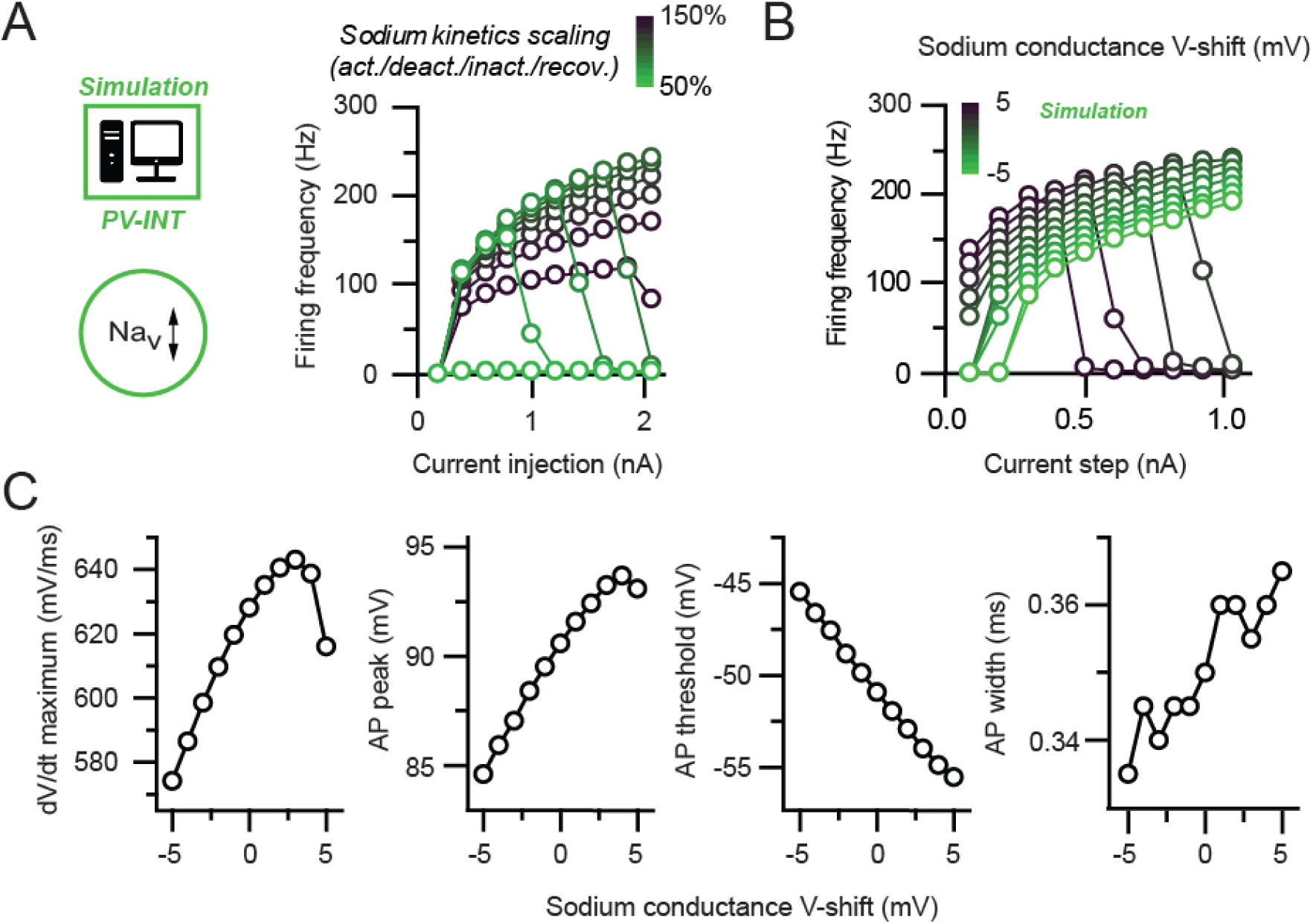
Na_v_ channel changes do not explain changes in PV interneuron excitability in 5xFAD mice. (A) Depiction of PV cell single compartmental model with modified Na_v_ channel properties. The relationship of injected current magnitude and AP firing frequency with varying Na_v_ kinetics is summarized. All four kinetic properties (activation, deactivation, inactivation, and recovery from inactivation) were simultaneously scaled together (± 50% of control) in the simulation. Near-threshold dampening of AP firing was observed with increased kinetics however this was accompanied by an overall reduction in AP firing rate at higher current injections. (B) Summary data showing the relationship of injected current magnitude and AP frequency following shifts in Na_v_ activation voltage (± 5mV from the control). Near-threshold dampening of AP firing was achieved through a hyperpolarizing shift however this was accompanied by a reduction in firing across all current injections, which was not observed in recordings from 5xFAD mice. (C) Additional datasets depicting the effect of shifting Na_v_ activation voltage on AP properties. Modifying activation voltage influenced all parameters including dV/dt maximum and AP threshold, which were not affected in recordings from 5xFAD mice.

**Figure S3, related to Figure 7.**
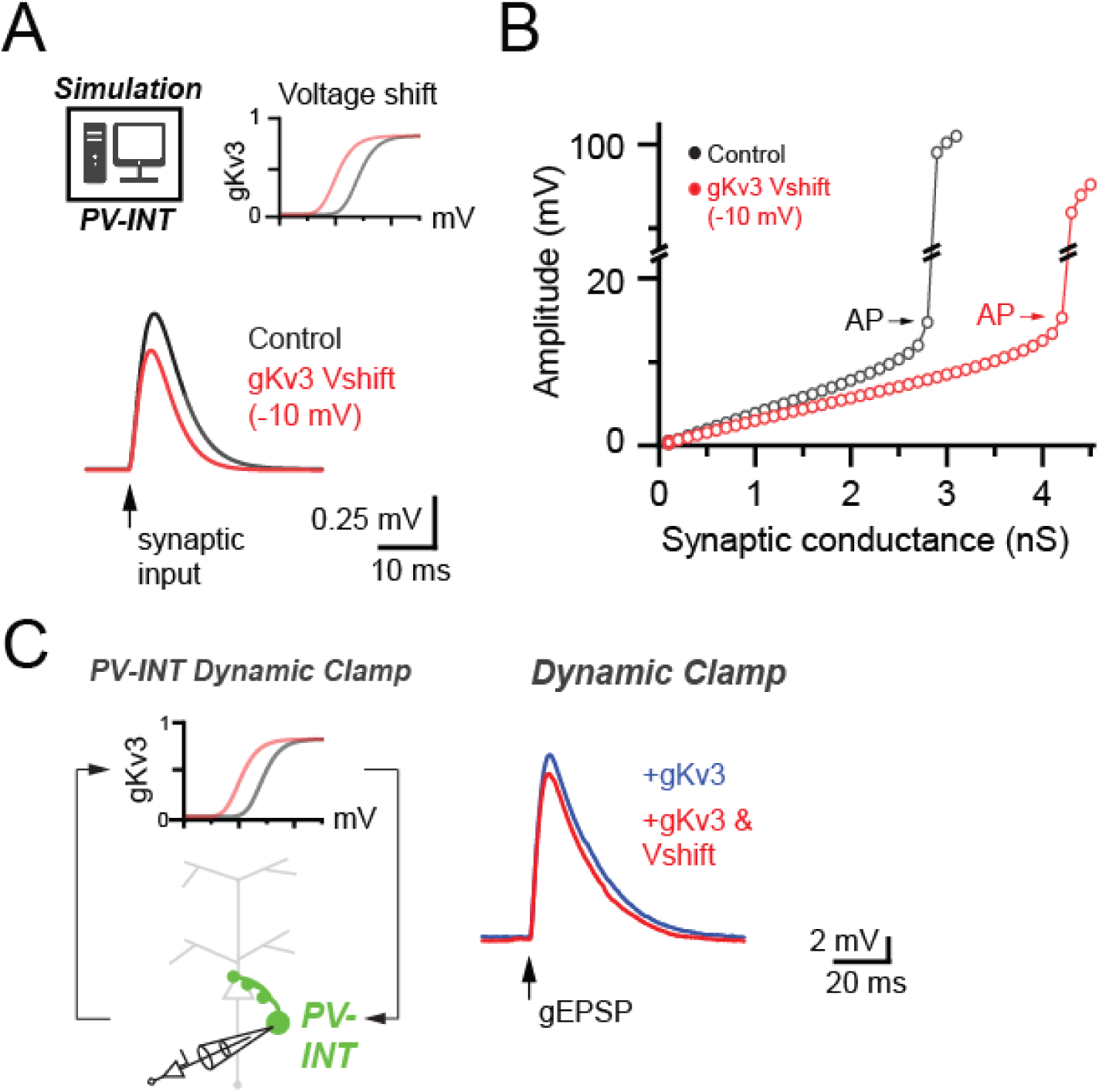
Effect of 5xFAD-related K_v_3 channel modulation on synaptically evoked subthreshold events. (A) Average subthreshold response of PV cell compartmental model following excitatory input in control conditions and with a relative −10 mV Vshift in K_v_3 activation voltage. (B) Differences in subthreshold excitatory event amplitudes evoked by increasing synaptic conductances. Simulation was run in control conditions and following leftward K_v_3 activation voltage shift. (C) Subthreshold gEPSPs evoked in dynamic clamp recordings from AAV.E2.GFP^+^ neurons. gEPSP conductance was calibrated such that stimuli reliably resulted in subthreshold events. For each condition during recordings, ∼10 time-locked gEPSP waveforms were averaged. Comparison of EPSP in +gK_v_3 (20 nS) and +gKv3 (20nS) with −10 mV Vshift are shown. −10mV K_v_3 relative Vshift was sufficient to reduce gEPSP charge (+gK_v_3, 252.2 ± 38.4 pC.; +gK_v_3 & Vshift, 221.9 ± 32.7 pC; p = 0.01, paired t-test; n = 5).

**Figure S4, related to Figure 8.**
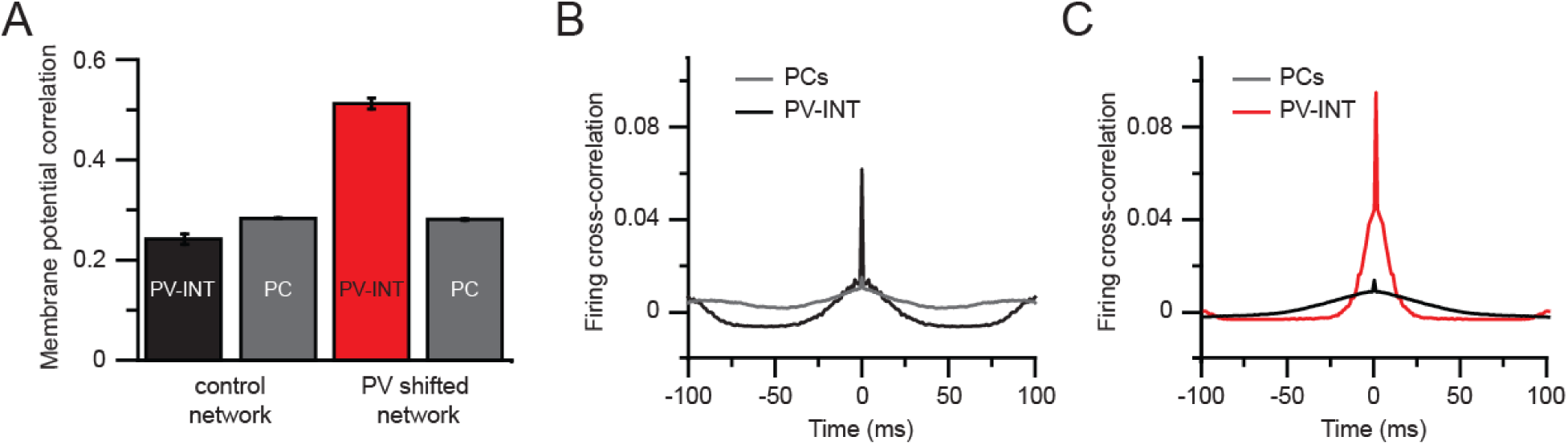
Circuit synchronization following PV-specific Kv3 modulation. (A) Membrane potential correlations within cell groups in control and −10 mV shifted gK_v_3 conditions. Correlations were measured as Pearson correlations coefficient comparing each individual cell. Data are expressed as mean (± SEM). (B) Firing cross-correlation of PV cells and PCs in a 200 ms time window. (C) Same as for panel B but in case of a network with −10 mV shifted gKv3 in PV cells.

## Methods

### Acute slice preparation

All animal procedures were approved by the Emory University IACUC. Acute slices from cortex were prepared from mature 5xFAD or littermate control (C57Bl/6J) mice (7-8 weeks old). Male and female 5xFAD mice and wild-type littermates were used for all experiments with data collected from ≥ 3 mice per experimental condition. Mice were first anesthetized and perfused with ice-cold cutting solution (in mM) 87 NaCl, 25 NaHO3, 2.5 KCl, 1.25 NaH2PO4, 7 MgCl2, 0.5 CaCl2, 10 glucose, and 7 sucrose. Thereafter, mice were killed by decapitation and the brain immediately removed by dissection. Brain slices (300 μm) were sectioned in the coronal plane using a vibrating blade microtome (VT1200S, Leica Biosystems) in the same solution. Slices were transferred to an incubation chamber and maintained at 34°C for ∼30 min and then at 23-24°C thereafter. During whole-cell recordings, slices were continuously perfused with (in mM) 128 NaCl, 26.2 NaHO3, 2.5 KCl, 1 NaH2PO4, 1.5 CaCl2, 1.5MgCl2 and 11 glucose, maintained at 30.0±0.5°C. All solutions were equilibrated and maintained with carbogen gas (95% O2/5% CO2) throughout.

### Electrophysiology

PV neurons were targeted for somatic whole-cell recording in layer 5 region of somatosensory cortex by combining gradient-contrast video-microscopy with epifluorescent illumination on custom-built or commercial (Olympus) upright microscopes. Electrophysiological recordings were obtained using Multiclamp 700B amplifiers (Molecular Devices). Signals were filtered at 6-10 kHz and sampled at 50 kHz with the Digidata 1440B digitizer (Molecular Devices). For whole cell recordings, borosilicate patch pipettes were filled with an intracellular solution containing (in mM) 124 potassium gluconate, 2 KCl, 9 HEPES, 4 MgCl2, 4 NaATP, 3 L-Ascorbic Acid and 0.5 NaGTP. Pipette capacitance was neutralized in all recordings and electrode series resistance compensated using bridge balance in current clamp. Liquid junction potentials were uncorrected. Recordings had a series resistance > 20 MΩ. Membrane potentials maintained near −70 mV (−70.7 ± 1.2 and −71.3 ± 0.8 mV; wild type and 5xFAD, respectively) during current clamp recordings using constant current bias. Action potential trains were initiated by somatic current injection (300 ms) normalized to the cellular capacitance in each recording measured immediately in voltage clamp after breakthrough (Taylor, 2012) (46.9 ± 2.5 and 46.3 ± 2.9 pF, n = 21 and 19, wild type and 5xFAD, respectively; p = 0.89; unpaired t-test. For quantification of individual AP parameters, the 1^st^ AP in a spike train at was analyzed at 9pA/pF for all cells. K^+^ channel activation curves were calculated as described (Rowan et al. 2016) using chord conductance (g) values from current peaks, and fit with a Boltzmann function. Activation time constants were obtained by fitting the rising phase of the K^+^ current with a single exponential function.

### Intracranial viral injections

Mice were injected with AAV(PHP.eB).E2.GFP in the SBFI vibrissal region of cortex. When performing viral injections, mice were head-fixed in a stereotactic platform (David Kopf Instruments) using ear bars, while under isoflurane anesthesia (1.8 - 2.2%). Thermoregulation was provided by a heating plate using a rectal thermocouple for biofeedback, thus maintaining core body temperature near 37°C. Bupivacaine was subcutaneously injected into the scalp to induce local anesthesia. A small incision was opened 5-10 minutes thereafter and a craniotomy was cut in the skull (< 0.5 μm in diameter) to allow access for the glass microinjection pipette. Coordinates (in mm from Bregma) for microinjection were X = ± 3.10 −3.50; Y = −2.1; α = 0°; Z = 0.85 – 0.95. Viral solution (titer 1×10^09^ to 1×10^12^ vg/mL) was injected slowly (∼0.02 μL min-1) by using a Picospritzer (0.3 μL total). After ejection of virus, the micropipette was held in place (5 min) before withdrawal. The scalp was closed with surgical sutures and Vetbond (3M) tissue adhesive and the animal was allowed to recover under analgesia provided by injection of carprofen and buprenorphine SR. After allowing for onset of expression, animals were sacrificed acute slices were harvested.

### Retro-orbital (RO) injection

Male and female Mice were given AAV retro-orbital injections as previously described in Chan et al. 2017. Mice were anesthetized with 1.8-2% isoflurane. AAV(PHP.eB).E2.GFP virus was titrated to 1×10^11^ vector genomes total and injected in C57B6/J mice to label of putative PV interneurons throughout cortex. As a control, PV-Cre mice (Jackson Laboratory; stock no. 008069); were injected with AAV(PHP.eB).Flex.tdTom (Addgene). Titrated virus was injected into the retro-orbital sinus of the left eye with a 31G x 5/16 TW needle on a 3/10 mL insulin syringe. Mice were kept on a heating pad for the duration of the procedure until recovery and then returned to their home cage for 2-3 weeks until sample collection.

### Fluorescent cell picking and qPCR

Manual cell picking was performed for single cell isolation. 12 mice (2 genotypes x 6 animals/group) were used for cell picking experiments. Acute slices (300 μm) were acquired from 5xFAD mice and their wild-type littermates at 7-8 weeks of age. Acute slices obtained as described above. Slices containing SBFI cortex were placed into cutting solution with 0.5 mg/mL protease (P5147–100MG, Sigma-Aldrich) for 60-minutes with continuous carbogen gas bubbling. Immediately after, slices were returned to room-temperature cutting solution for 10 minutes. Slices were then micro-dissected to isolate the cortical region containing GFP^+^ or tdTom expressing cells using an epi-fluorescent stereoscope (Olympus SZX12). Samples were then manually triturated in cutting solution with 1% Fetal Bovine Serum (F2442–50 mL, Sigma Aldrich) into a single-cell suspension. The sample was then diluted with ∼300 uL of cutting solution, dropped onto a Sylgard (DOW) coated petri dish, and cells were allowed 10 minutes to settle. The remainder of the dish was then filled with pre-bubbled cutting solution. Cells were selected using epi-fluorescent illumination under an inverted microscope (Olympus IX71) using a pulled borosilicate glass pipette connected to a filter-tipped stopcock. ∼200 picked cells were stored in RLT buffer (Cat. No. 79216 – 220 mL, Qiagen) with 1% 2-Mercaptoethanol (M6250-100ML, Sigma-Aldrich) at −80°C until cDNA isolation. cDNA was generated from each sample using an RNAseq library prep method. A cDNA library was created with the CellAmp™ Whole Transcriptome Amplification Kit (#3734, Takara Bio) to allow for real-time PCR (qPCR) to be conducted. qPCR was then conducted with the following primers:

GAPDH (Mm99999915_g1, Taqman), PVALB (Mm.2766, Taqman), SCN1A

(Mm00450580_m1, Taqman), SCN8A (Mm00488119_m1, Taqman), KCNC1

(Mm00657708_m1, Taqman), KCNC2 (Mm01234232_m1, Taqman), KCNC3

(Mm00434614_m1, Taqman), KCNC4 (Mm00521443_m1, Taqman), GRIN1

(Mm00433790_m1, Taqman), GRIN2A (Mm00433802_m1, Taqman), GRIN2B

(Mm00433820_m1, Taqman).

Results of qPCR were analyzed using the Common Base Method with expression normalized to GAPDH. ΔCt values were averaged between triplicate samples from each mouse.

### PC cell NEURON modeling

Computer simulations were performed using the NEURON simulation environment (version 7.5 and 7.6, downloaded from http://neuron.yale.edu). For PV interneuron models a single 20µm × 20µm compartment was created and was equipped by sodium, potassium and leak conductances. The passive background of the cell was adjusted to recreate passive membrane potential responses of whole-cell recorded PV INs for given stimulus intensities. The sodium conductance was based on the built-in Hodgkin-Huxley model of NEURON with freely adjustable sets of parameters (Oláh et al., 2021). The PV potassium conductance was implemented based on a previous publication (Lien and Jonas, 2003) constrained by our outside-out patch recordings. The steady state activation was governed by the following equation:

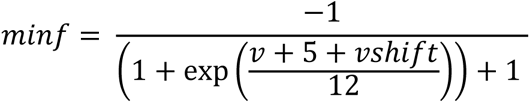

where v is local membrane potential and *VShift* is the applied voltage shift in order to adjust membrane potential dependence. The steady state inactivation was set as follows:

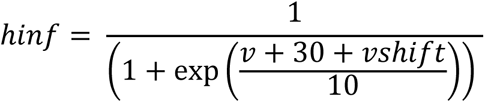

The activation and deactivation time constant was defined as:

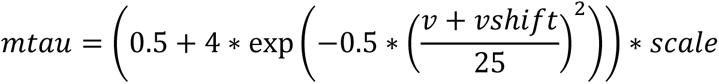

where scale was the parameter by which kinetics were adjusted. Inactivation time constant was set to 1000 ms. Synaptic inputs for examining firing responses under more naturalistic network conditions were supplemented by using NEURON’s built-in AlphaSynapse class. During the simulation (1 second), 1000 individual excitatory synapses and 500 inhibitory synapses were added with random timing, 10 nS synaptic conductance, and 0 or −90 mV reversal potential, respectively.

### Network simulations

Network simulations were carried out with the class representation of the previously detailed FS model cell, and a newly constructed pyramidal cell (PC) mode, which was a slight modification of a bursting model cell described by earlier (Pospischil et al., 2008). 200 PC and 20 PV cells were used and connected with accordance to previous publications. Recurrent PC connectivity was set to 10% (Markram et al., 2015), PV-to-PC connectivity was set to 36% (Packer and Yuste, 2011), PV cell recurrent connections occurred with 78% probability, and gap junction connectivity between these cells was 61% (Galarreta and Hestrin, 2002). Finally, PC innervated PV cells with 80% chance (Hofer et al., 2011). All simulated cells received constant current injections in order to elicit baseline firing at variable frequencies. The network construction was done in several consecutive steps. First, PV cells were connected to each other with chemical synapses constrained to elicit moderate network synchronization (Wang and Buzsáki, 1996). Next, PV cells were connected with gap junctions, were gap junction conductance was set to a value, which could synchronize the network further. PV cells inhibited PC cells with less inputs less than 1 mV in amplitude (Packer Yuste 2011), similarly to PC to PV connections (Hofer et al. 2011). Firing correlations and power spectrum was analyzed in Python. All modelling related code will be made available upon request.

### Dynamic clamp

The dynamic clamp system was built in-house based on a previous publication (Desai Johnston 2017), related online available materials (www.dynamicclamp.com). The equations governing the implemented gKdr were identical to those used in the NEURON model construction. Synaptic conductances were built-in predefined conductances available from www.dynamicclamp.com.

### Stats and Analysis

Custom python scripts, Axograph, Graphpad Prism (Graphpad Software), and Excel (Microsoft) were used for analysis with values in text and figures. Statistical differences were deemed significant with α values of p < 0.05. Unpaired and paired t-tests were used for unmatched and matched parametric datasets, respectively. Where appropriate, group data were compared with 1 or 2-way ANOVA and significance between groups noted in figures was determined with Tukey’s or Sidak’s multiple post-hoc comparison tests. Normality was determined using D’Agostino & Pearson omnibus or Shapiro-Wilk tests.

## Notes

### Competing Interest Statement

The authors have declared no competing interest.

